# The *Toxoplasma* centrocone houses cell cycle regulatory factors

**DOI:** 10.1101/122465

**Authors:** Anatoli Naumov, Stella Kratzer, Li-Min Ting, Kami Kim, Elena S. Suvorova, Michael W. White

## Abstract

Our knowledge of cell cycle regulatory mechanisms in apicomplexan parasites is very limited. In this study, we describe a novel *Toxoplasma gondii* factor, essential for chromosome replication 1 (ECR1), that has a vital role in chromosome replication and the regulation of cytoplasmic and nuclear mitotic structures. ECR1 was discovered by complementation of a temperature sensitive (ts) mutant that suffers lethal, uncontrolled chromosome replication at 40^°^C similar to a ts-mutant carrying a defect in topoisomerase. ECR1 is a 52kDa protein containing divergent RING and TRAF-Sina like zinc-binding domains that is dynamically expressed in the tachyzoite cell cycle. ECR1 first appears in the centrocone compartment of the nuclear envelope in early S phase and then in the nucleus in late S phase where it reaches maximum expression. Following nuclear division, but before daughters resolve from the mother, ECR1 is down regulated and is absent in new daughter parasites. The proteomics of ECR1 identified interactions with the ubiquitin-mediated protein degradation machinery and the minichromosome maintenance complex and the loss of ECR1 led to increased stability of a key member of this complex, MCM2. ECR1 also forms a stable complex with the CDK-related kinase, TgCrk5, which shares a similar cell cycle expression and localization during tachyzoite replication. Altogether, the results of this study suggest ECR1 may be a unique E3 ligase that regulates DNA licensing and other mitotic processes. Importantly, the localization of ECR1/TgCrk5 in the centrocone indicates this Apicomplexa-specific spindle compartment houses important regulatory factors that control the parasite cell cycle.

**IMPORTANCE:** Parasites of the apicomplexan family are important causes of human disease including malaria, toxoplasmosis, and cryptosporidiosis. Parasite growth is the underlying cause of pathogenesis, yet despite this importance the molecular basis for parasite replication is poorly understood. Filling this knowledge gap cannot be accomplished by mining recent whole genome sequencing because apicomplexan cell cycles differ substantially and lack many of the key regulatory factors of well-studied yeast and mammalian cell division models. We have utilized forward genetics to discover essential factors that regulate cell division in these parasites using the *Toxoplasma gondii* model. An example of this approach is described here with the discovery of a putative E3 ligase/protein kinase mechanism involved in regulating chromosome replication and mitotic processes of asexual stage parasites.

## INTRODUCTION

Highly efficient asexual replication is fundamental to the ability of apicomplexan parasites to spread infections in their hosts and this is evident by the action of drugs used to combat these infections as the best treatments all reduce or block parasite proliferation. Unfortunately, existing therapies, particularly against malaria (1), are under constant pressure from acquired parasite drug resistance requiring a continuing search for new treatments. The peculiar proliferative cell cycles of Apicomplexa parasites differ substantially from the hosts they inhabit and it seems safe to predict that a better understanding of the molecular basis of parasite cell division could yield new drug targets. For most apicomplexan species replication is characterized by a sequence of two chromosome replication cycles (2). A single G1 phase that is roughly proportional in length to the number of parasites produced precedes a first chromosome cycle (S/M_n_-nuclear cycle) whose biosynthetic focus is genome replication followed by a single chromosome cycle (S/M_n+1_-budding cycle) that produces infectious parasites assembled internally (endopolygeny) or adjacent the plasma lemma (schizogony) of the mother parasite (2). How genome fidelity is preserved through variable rounds of apicomplexan chromosome replication is not understood as many known regulators of cell cycle progression in yeast and multicellular eukaryotes are missing in these parasites (3). Further, the basic checkpoint mechanisms that regulate the cell cycle transitions in Apicomplexa replication are also poorly understood. It is generally not known what molecular mechanisms control G1 to S phase commitment, S phase progression, chromosome segregation or what regulatory factors enable apicomplexan parasites to suspend budding until the last round of chromosome replication.

Analysis of many Apicomplexa genomes shows that genome mining by itself will not yield the molecular basis of cell cycle control in these parasites, which will require bench discoveries. To this end, the *T. gondii* tachyzoite has provided a valuable model to study cell cycle mechanisms. The *T. gondii* tachyzoite utilizes an abbreviated cell cycle (endodyogeny) to produce two progeny from each mother cell (4). The phases of the tachyzoite cell cycle are defined (5, 6) and feature G1, S, and overlapping mitotic and cytokinetic processes that can be now distinguished with specific protein markers (2). Key cell cycle transitions where checkpoint mechanisms likely operate have also been revealed by the development of synchrony methods (6) and the study of cell cycle ts-mutants (2, 7, 8). At a minimum, *T. gondii* tachyzoites possess mechanisms that control G1 progression, commitment to chromosome replication, and may regulate a short G2 phase that is characterized by an unusual 1.8N genome content (3). The discovery of a bipartite centrosome with independent cores regulating karyokinesis or cytokinesis indicate additional points of cell cycle control (2). These previous studies have paved the way for understanding the regulatory basis of cell division in these parasites. In this study, we begin to fill in these details with the discovery of a unique protein complex composed of a dual zinc finger protein and protein kinase belonging to the cyclin-dependent family (<u>C</u>DK-<u>r</u>elated <u>k</u>inases=<u>C</u>rk) that is involved in regulating chromosome replication in the tachyzoite. This protein complex is initially housed in the unique spindle compartment of the Apicomplexa called the centrocone where it may regulate the connection between nuclear and cytoplasmic mitotic structures that must interact in order to coordinate the complex events associated with the tachyzoite mitosis.

## RESULTS

### *Toxoplasma* chromosome replication and segregation requires a conserved topoisomerase and a novel dual zinc finger protein

The peculiar cell cycles of the Apicomplexa suggest that molecular controls of replication in these parasites may be equally unique. To this end, we have used forward genetics to identify essential cell cycle factors in *T. gondii* (2, 7-11). The temperature sensitive (ts) mutants 11-51A1 and 13-64D5 from our collection of ∼150 growth mutants (7) share similar lethal temperature defects (Fig. 1A, grow curves). When shifted to 40^°^C, both ts-mutant parasites arrested quickly (within 0-2 divisions) and this was associated with the loss of near-diploid to diploid genomic DNA content and the increase of daughter parasites possessing 1N and sub-1N contents as analyzed by flow cytometry (Fig. 1B). Cell division of each ts-mutant strain at 34^°^C was similar to the parent (Fig. 1A and 1C), while at 40^°^C ts-mutants 11-51A1 and 13-64D5 showed severe defects in mitotic and cytokinetic processes (Fig. 1C) characterized by loss of normal nuclei and daughter parasite stoichiometry (normal ratios, 1:2 & 1:1). Daughter buds formed that often had plastid DNA, but many buds lacked nuclear DNA (Fig. 1C) indicating that cytokinesis and karyokinesis was uncoupled in both ts-mutants. By immunofluorescent assay (IFA) masses of unpackaged nuclear DNA accumulated in the residual mother parasite (Fig. 1C) indicating that restrictions on chromosome replication may also be lost at high temperature. The accumulation of ts-mutant parasites with sub-1N DNA contents observed in flow cytometry analysis (Fig. 1B) is consistent with the inability of these parasites to package their DNA into daughter parasites.

**FIG. 1.**
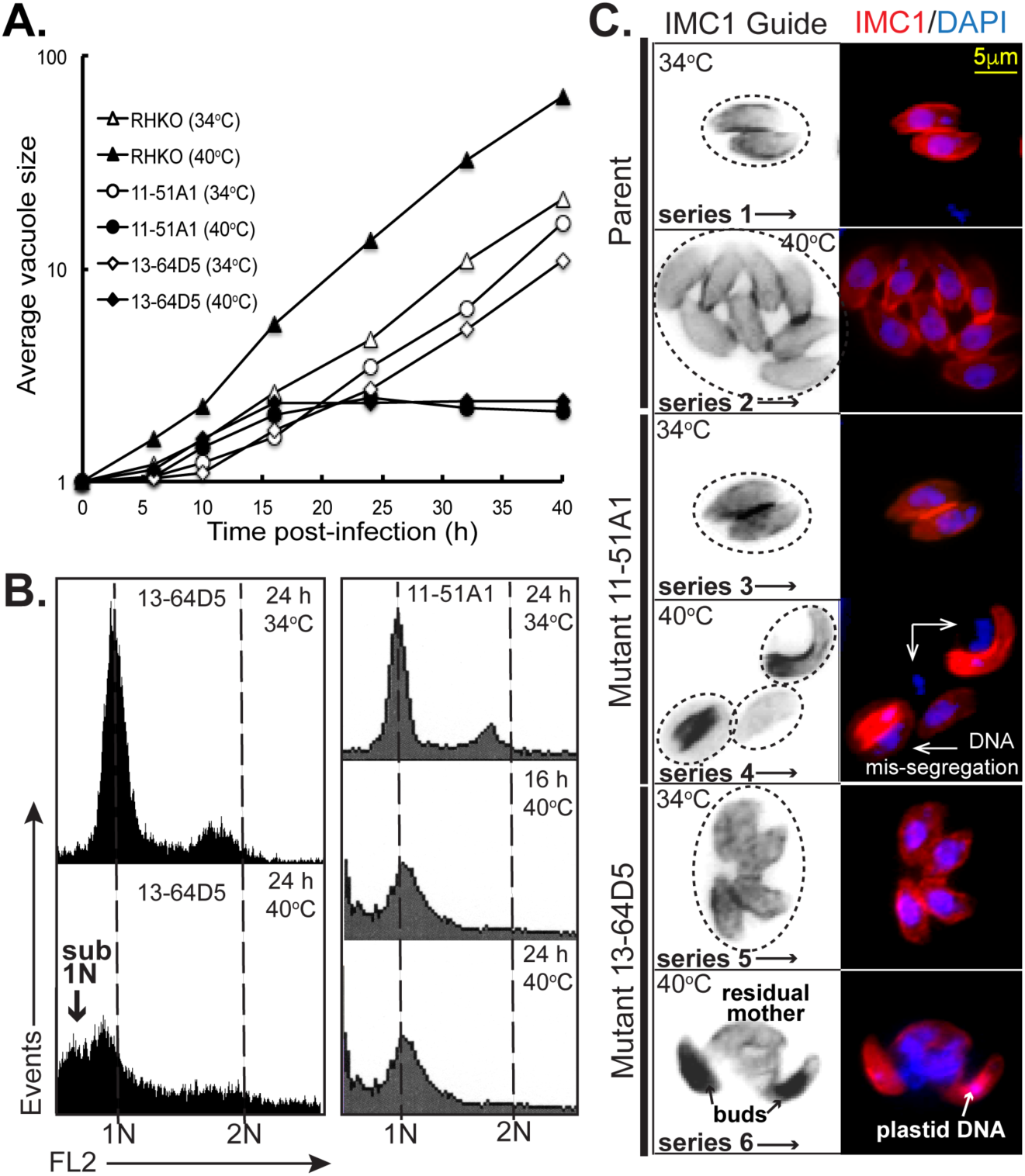
Temperature-sensitive ts-mutants defective in chromosome replication. **[A.]** The growth of chemical ts-mutants 13-64D5 and 11-51A1 was inhibited by high temperature leading to lethal arrest. At the nonpermissive temperature of 40^°^C, the ts-mutants arrested within 1-2 divisions, whereas at the permissive temperature of 34^°^C, the ts-mutants replicated similar to the parent strain. Note that all of the strains, including the parent strain, grew more slowly at 34^°^C as compared to the parent strain at 40^°^C**. [B.]** Flow cytometric analysis of EtOH fixed ts-mutant parasites stained with propidium iodide after RNAase treatment showed similar sub-1N aneuploidy that is consistent with chromosome loss at 40^°^C. The DNA content of the ts-mutants grown at 34^°^C exhibits the 1N and 1.8N peaks representative of an asynchronous growing tachyzoite population (6). For each sample, 10,000 parasites were analyzed (y axis) using an FL-2 linear scale; 1N and 2N DNA fluorescent values are indicated on the x-axis and in the vertical dashed lines. **[C.]** IFA analysis of ts-mutants 13-64D5 and 11-51A1 compared to the parent strain grown at 34^°^C (image series 1-5) and 40^°^C (image series 2-6). The parasites were stained for DNA with DAPI and internal daughter budding with αIMC1. The IMC1 Guide is a non-colored/inverted image in order to better demark individual mother parasites and daughter buds. Circles in the IMC1 Guide indicate the number of vacuoles and the approximate size base on DIC images (results not shown). Mutant images at 40^°^C (series 4 and 6) show a similar phenotype of unequal masses of chromosomal DNA accumulated free of abnormal daughter parasites. Note that the small DAPI focus in the buds of series 6 is plastid DNA. Scale bar in the series 6 fluorescent image applies to all images.

To identify the defective gene in ts-mutant 11-51A1, parasites were genetically complemented with a *T. gondii* cosmid-based genomic library under 40^°^C and pyrimethamine selection to ensure rescued parasites possessed cosmid DNA (7). Marker-rescue of the resulting transgenic parasites (without cloning) identified a locus on chromosome VIIb as the location of the ts-mutation (Fig. 2A). Complementation of the original ts-mutant 11-51A1 with PCR fragments spanning each of the four central genes in the locus determined gene 4 (TGME49_258790) was the defective gene. Sequencing of ts-mutant gene 4 pinpointed a non-synonymous mutation that changed a leucine to histidine at residue 413 as the cause of temperature sensitivity in this chemical ts-mutant. Gene 4 encodes a ∼52kDa protein containing divergent RING (PF13923, e-08) and TRAF-like Sina (e-09) zinc binding domains (Fig. 2A). Due to the requirement of this protein for proper tachyzoite chromosome replication and segregation, gene 4 was named **E**ssential for **C**hromosome **R**eplication **1** (ECR1). Western analysis of ectopically expressed ECR1^Myc^ and tsECR1^Myc^ revealed that protein instability at high temperature (Fig. S1A) was likely responsible for the loss of tsECR1 function and the underlying cause of temperature lethality of the original 11-51A1 ts-mutant strain.

**FIG. 2.**
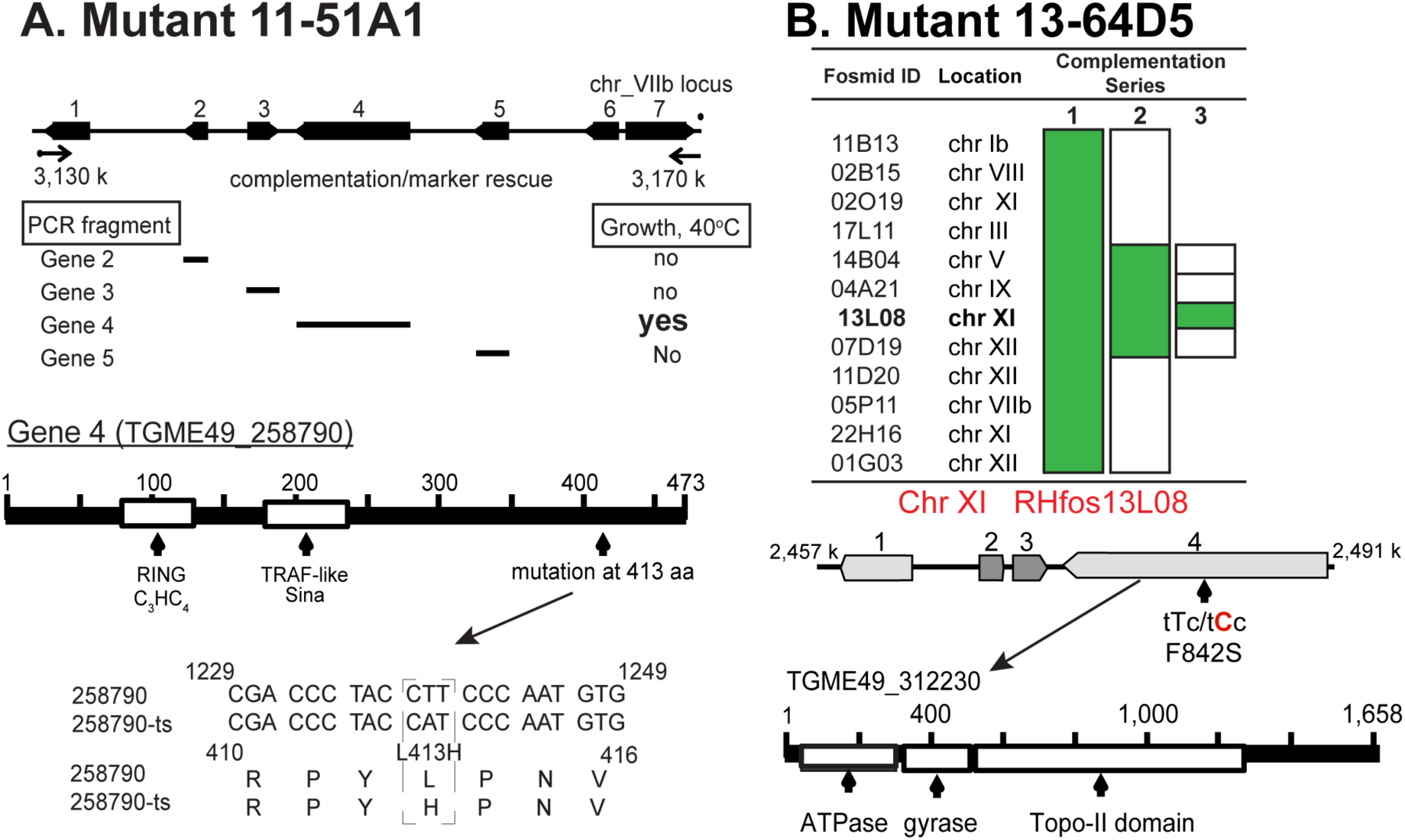
Identification of the defective genes in ts-mutants 13-64D5 and 11-51A1. **[A.]** Genetic complementation of ts-mutant 11-51A1 with cosmid genomic libraries followed by marker-rescue identified the defective locus to chromosome VIIb (see top diagram). The locus was further resolved using PCR products spanning each of the central four genes. Only gene 4 (TGME49_258790) rescued ts-mutant 11-51A1 at 40^°^C and was sequenced to identify the ts-mutation. The mutation and size and major protein domains for Gene 4 are shown**. [B.]** The genome of ts-mutant 13-64D5 was sequenced to >150-fold coverage identifying 16 non-synonymous mutations. Fosmid clones from a mapped *T. gondii* genetic library that spanned 12 of the sequenced mutations were combined and used to successfully complement ts-mutant 13-64D5 (complementation series 1). Genetic complementation with fosmid clones re-mixed into three groups (series 2) followed by complementation of four individual fosmids (series 3) identified a region on chromosome XI as the locus carrying the mutation responsible for high temperature sensitivity. In the identified XI locus, ts-mutant 13-64D5 harbors a single mutation (F842S) in a *T. gondii* ortholog of topoisomerase II (TgTopo-II) that is rescued by parental copy of TgTopo-II on fosmid RHfos13L08.

In order to identify the defective gene in ts-mutant 13-64D5, we used a whole-genome sequencing/complementation approach as previously described (12). Briefly, ts-mutant 13-64D5 genomic DNA was sequenced to >150x coverage on the Illumina HiSeq 2500 platform and non-synonymous mutations were identified in 16 genes. Three of these genes were eliminated due to the lack of tachyzoite expression and/or a conserved amino acid substitution. Fosmid clone DNA carrying *T. gondii* genomic inserts (12) encompassing the remaining 13 genes (fosmid 02O19 spanned two SNPs) were purified and combined into a single mix for genetic complementation (Fig. 2B, complementation series 1). Successful rescue of ts-mutant 13-64D5 with the combined 12 fosmid DNAs was followed by complementation with the 12 fosmid DNAs remixed in 3 groups of 4 (Fig. 2, series 2), and finally, four individual fosmid clones were tested that identified a single fosmid clone able to rescue high temperature sensitivity (series 3). The sole ts-mutant SNP in the genomic locus spanned by the fosmid clone RHfos13L08 insert is a T<u>T</u>C to T<u>C</u>C change that leads to serine substituting phenylalanine at amino acid residue 842 in the *T. gondii* ortholog of eukaryotic topoisomerase II (TGME49_312230). *T. gondii* topoisomerase II (TgTopo-II) is a 185 kDa protein with N-terminal ATPase and gyrase domains, central topoisomerase domain, and unique C-terminal tail (Fig. 2B).

### TgTopo-II and ECR1 are cell cycle regulated nuclear factors

TgTopo-II and ECR1 proteins represent distinct evolutionary histories in apicomplexan parasites (Fig. S2). Topo-II is highly conserved in eukaryotes and is present in all apicomplexan species sequenced (Fig. S2A). By contrast, ECR1 is limited to modern coccidian species (Fig. S2B). However, we found highly divergent ECR1 orthologs in ancestral free-living protozoan *Chromera velia* and, unexpectedly, in filamentous fungi (Fig. S2A), indicating that the last common ancestor of eukaryotes might have encoded ECR1 related factor. Marginal sequence similarity covers only functional regions of these hypothetical proteins, namely a RING and a TRAF-like Sina (<u>s</u>even <u>i</u>n <u>a</u>bsentia) domain (Fig. S2B). The fact that Sina factors in higher eukaryotes provide a link between DNA damage and β-catenin degradation suggests the ancestral role for ECR1-related factors in sensing the DNA state (13).

TgTopo-II and ECR1 mRNAs are similarly cell cycle regulated with peak expression in the tachyzoite S phase (Fig. 3A). To determine if the profile of the encoded proteins was also periodic, each protein was C-terminally fused to 3 copies of the HA epitope by genetic knock-in in order to preserve native expression. TgTopo-II^HA^ was exclusively nuclear and expressed throughout the tachyzoite cell cycle with an increase in expression during S phase (Fig. 3B) similar to the encoded mRNA (Fig. 3A). TgTopo-II^HA^ co-localized with nuclear DNA (Fig. 3B, see DAPI/αHA co-staining) and during telophase this factor was concentrated at the tips of u-shaped nuclei beginning to divide (Fig. 3B, inset image). Like TgTopo-II^HA^, ECR1 tagged by genetic knock-in was determined to be a nuclear factor, although the expression of ECR1^HA^ was much more dynamic. ECR1^HA^ first appeared following duplication of the centrosome outer core (see example Fig. 4, images 1-3) concentrated in the apicomplexan-specific centrocone prior to the duplication of this structure (Fig. 3C, see guide reference and MORN1 co-stain)(2). Following duplication and separation of the centrocone (early mitosis) ECR1^HA^ became exclusively nuclear where it reached maximum levels of expression. ECR1^HA^ was down regulated during budding (Fig. 3C, S/M images) and was absent from G1 parasites (Fig. 3C, G1 images). Altogether, these results indicate that ECR1 is expressed starting in early S phase through early mitosis.

**FIG. 3.**
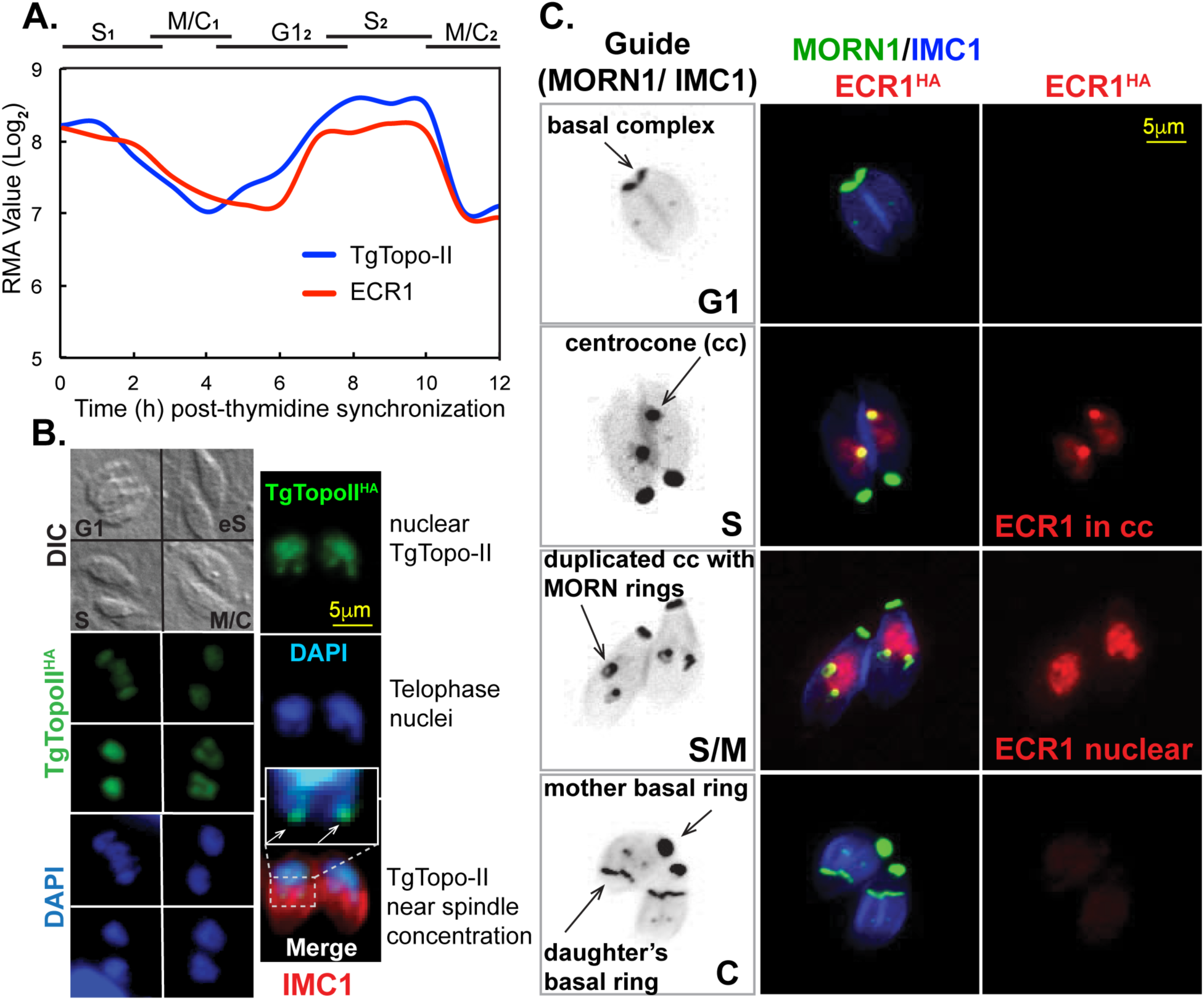
ECR1 and TgTopo-II are cell cycle regulated nuclear factors. **[A.]** The mRNAs encoding ECR1 and TgTopo-II are similarly cell cycle regulated with peak expression in S phase consistent with potential roles in DNA replication. **[B. and C.]** The wild type genes ECR1 (defective in ts-mutant 11-51A1) and TgTopo-II (defective in 13-64D5) were epitope tagged with 3xHA by genetic knock-in. IFA analysis of TgTopo-II^HA^ and ECR1^HA^ revealed a cell cycle expression profile matching their respective cyclical mRNA profiles. Both factors were maximally expressed during S phase before falling to low or undetectable levels in early G1 parasites. In **[B.],** TgTopo-II^HA^ transgenic parasites were co-stained with aHA (green), aIMC1 (red) and DAPI (blue) to visualize DNA. DIC image on top shows four vacuoles in different cell cycle phases as indicated. TgTopo-II^HA^ was exclusively nuclear throughout the cell cycle with some protein concentrating in the leading edge of nuclei (inset image of merged panel) beginning to undergo nuclear division (telophase). In **[C.],** ECR1^HA^ (red) transgenic parasites were costained for MORN1 (green) to visualize the centrocone and basal complexes and also for IMC1 to reveal the mother and daughter inner membrane complexes (blue). The cell cycle expression of ECR1^HA^ was complex; first appearance in the centrocone in early S, then exclusive nuclear in late S, and disappearing prior to nuclear division. Guide panels are non-colorized/inverted IMC1/MORN1 merged images labeled for centrocone (spindle pole) and basal complexes (front edge of growing daughter buds). The cell cycle phases are indicated in the lower right corner of the guide panels. Magnification scale bars indicated apply to all images in each series **[B. and C.].**

**FIG. 4.**
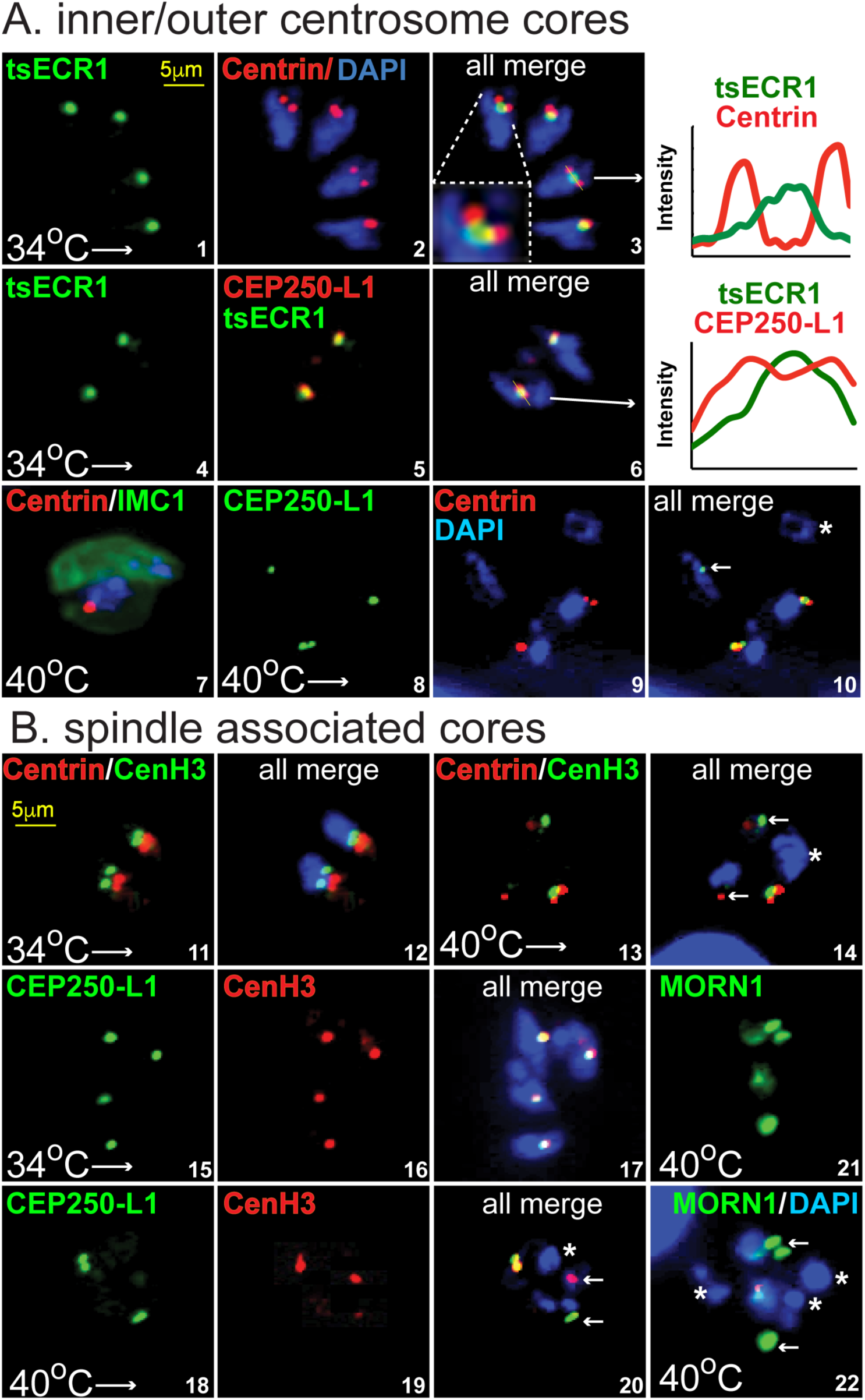
ECR1 is required for replicating and assembling key mitotic structures. Mitotic structures were examined in tsECR1^HA^ ts-mutant parasites (see Fig. S1B for construction) at permissive and nonpermissive temperatures. Visualization of the inner (CEP250-L1^HA^) and outer (acentrin) cores of the tachyzoite centrosome (2) and the centrocone (tsECR1^HA^ at 34^°^C) and kinetochore (aCenH3) was accomplished by epitope-tagging specific integral proteins (by genetic knock-in) or by staining with protein-specific antibodies as indicated. DAPI staining was used to visualize genomic DNA content and magnification scale bars are indicated once in each series**. [A.]** Centrosome structures (centrosome inner and outer cores) of growing tsECR1^Myc^ mutant parasites at 34^°^C (images 1-6) were compared to growth arrested parasites at 40^°^C (images 7-10). The following pairwise proteins were evaluated at 34^°^C: images 1-3, tsECR1^Myc^ in the centrocone (green) versus centrin expression (red) in the outer centrosome core; images 4-6, tsECR1^Myc^ (green) in the centrocone versus CEP250-L1^HA^ in the centrosome inner core (each red stains). The inset of image 3 and the graphs of fluorescent scans (lines through the structures indicated by arrows) in images 3 and 6 show tsECR^Myc^ localization in relation to centrin in the outer and CEP250-L1^HA^ in the inner centrosome cores. Note the clear physical resolution of tsECR1^Myc^ in the centrocone from centrin in the centrosome outer core. Pairwise protein expression at 40^°^C: image 7, centrin versus IMC1 in the tsECR1^Myc^ mutant strain (tsECR1^Myc^ is absent in these parasites at 40^°^C); images 8-10, CEP250-L1^HA^ versus centrin expression in the tsECR1^Myc^ transgenic strain. Note the abnormal nuclei/core stoichiometry and loss of alignment of the centrosome cores. Image 10, the nucleus indicated by a star has no centrosome cores, while the nucleus indicated by an arrow only has an inner core containing CEP250-L1^HA^**. [B.]** Spindle-associated structures (centrocone, inner core, and kinetochore) of tsECR1^Myc^ mutant parasites at 34^°^C (images 11,12,15-17) were compared to growth at 40^°^C (images 13,14, 18-22). Histone CenH3 of the kinetochore (green) was compared to centrin (red) of the outer centrosome core (images 11-14). Normal replication and inside/outside close alignment of these structures to the nucleus (blue, DAPI staining) was observed in tsECR1^Myc^ parasites grown at 34^°^C (images 11,12), whereas, the loss of tsECR1^Myc^ at 40^°^C lead to severe physical and stoichiometric changes. The star in image 14 marks a nucleus lacking both centrosome and kinetochore structures and the arrows indicate single, free kinetochore and outer core structures in the ts-mutant parasite cytoplasm. The inner core protein CEP250-L1^HA^ was compared to CenH3 in parasites grown at 34^°^C (images 15-17) versus 40^°^C (images 18-20). Note that the normal tight alignment of the inner centrosome core with the kinetochore is lost in tsECR1^Myc^ parasites grown at high temperature. Image 20, star marks a nucleus lacking inner and kinetochore structures, while the arrows mark single core structures of each type. Image 21 and 22, MORN1 versus DAPI staining in tsECR1^Myc^ parasites at 40^°^C possess nuclei (stars) without an intact centrocone.

### Disruption of ECR1 leads to severe morphological defects in mitotic structures

To further investigate ECR1 function, we introduced the L413H mutation along with a 3xMyc epitope tag by knock-in strategy into the ECR1 gene sequence of a parental strain not previously exposed to chemical mutagenesis (Fig. S1B, diagram). The conversion of native ECR1 into tsECR1^Myc^ generated a temperature sensitive strain with identical mitotic and cytokinetic defects at high temperature as the original 11-51A1 ts-mutant including the disruption of nuclei packaging into daughter buds (Fig. S1C and Fig. 4A, image 7). At the permissive temperature (34^°^C), the recreated tsECR1^Myc^ strain replicated similar to the parent strain and possessed healthy centrosome structures (Fig. S1C). Similar to the epitope-tagged native protein (Fig. 3C), tsECR1^Myc^ at 34^°^C was first detected in the centrocone following duplication of the centrosome outer core (Fig. 4A, images 1-3). In normal replicating tachyzoites the centrocone is closely aligned with the centrosome inner core (Fig. 4A, images 4-6) and resolved from the outer core of the centrosome (see ref. 2). By contrast, centrosome control structures were dramatically affected by the loss of tsECR1^Myc^ at 40^°^C (Fig. 4A and B). Inner (images 8-10) and outer (images 7,13,14) core physical and stoichiometric balance was disrupted leading to some nuclear packages lacking any associated centrosome core. The changes resulting from the loss of tsECR1^Myc^ at 40^°^C were not confined to centrosome defects. The usual close alignment of the intranuclear kinetochore (visualized by CenH3)(14) with the centrocone and inner centrosome core was uncoupled and the CenH3 structure became extranuclear (Fig. 4B). For many nuclear packages there was no evidence of CenH3 staining present in or nearby the distinct DAPI staining packages (images 14, 20). Finally, the most dramatic change was the nearly complete loss of MORN1 centrocone staining (images 21, 22) suggesting the loss of tsECR1^Myc^ at 40^°^C caused severe defects in the intranuclear organization of the mitotic machinery, including centrocone and associated kinetochores.

### The ECR1 interactome reveals functions in regulating the parasite cell cycle

In order to better understand the ECR1 mechanism, we performed IP/LC-MS/MS on ECR1^HA^ parasites (Fig. 5 and Dataset S1). Purified ECR1^HA^ complexes from whole cell lysates were resolved by electrophoresis and individual gel slices subjected to mass spectrometry analysis (see Material and Methods). This experiment identified 123 proteins including ECR1^HA^ itself (24.9% coverage, 47 total spectra) (Dataset S1). Importantly, many nuclear factors were co-immunoprecipitated with ECR1^HA^, including 5 of 6 subunits of the minichromosome maintenance complex (MCM_2-7_; 1.34%, 69 spectra), which is the essential helicase of semiconservative DNA replication in eukaryotes (15). The ECR1^HA^ pull down also revealed significant interactions with the ubiquitin-mediated protein degradation machinery (24 proteins) and with one (TgCrk5, TGME49_229020) of ten serine/threonine protein kinases encoded in the *T. gondii* genome that are distantly related to human cyclin-dependent kinases (16). Interaction of ECR1 with an E1 ubiquitin-activating enzyme, two E2 ubiquitin-conjugating enzymes, UBX domain containing protein, and components of the 26S proteasome (α and β, and regulatory subunits) indicate that the RING domain protein ECR1 may function as an E3 ubiquitin ligase (see Fig. 5).

**FIG. 5.**
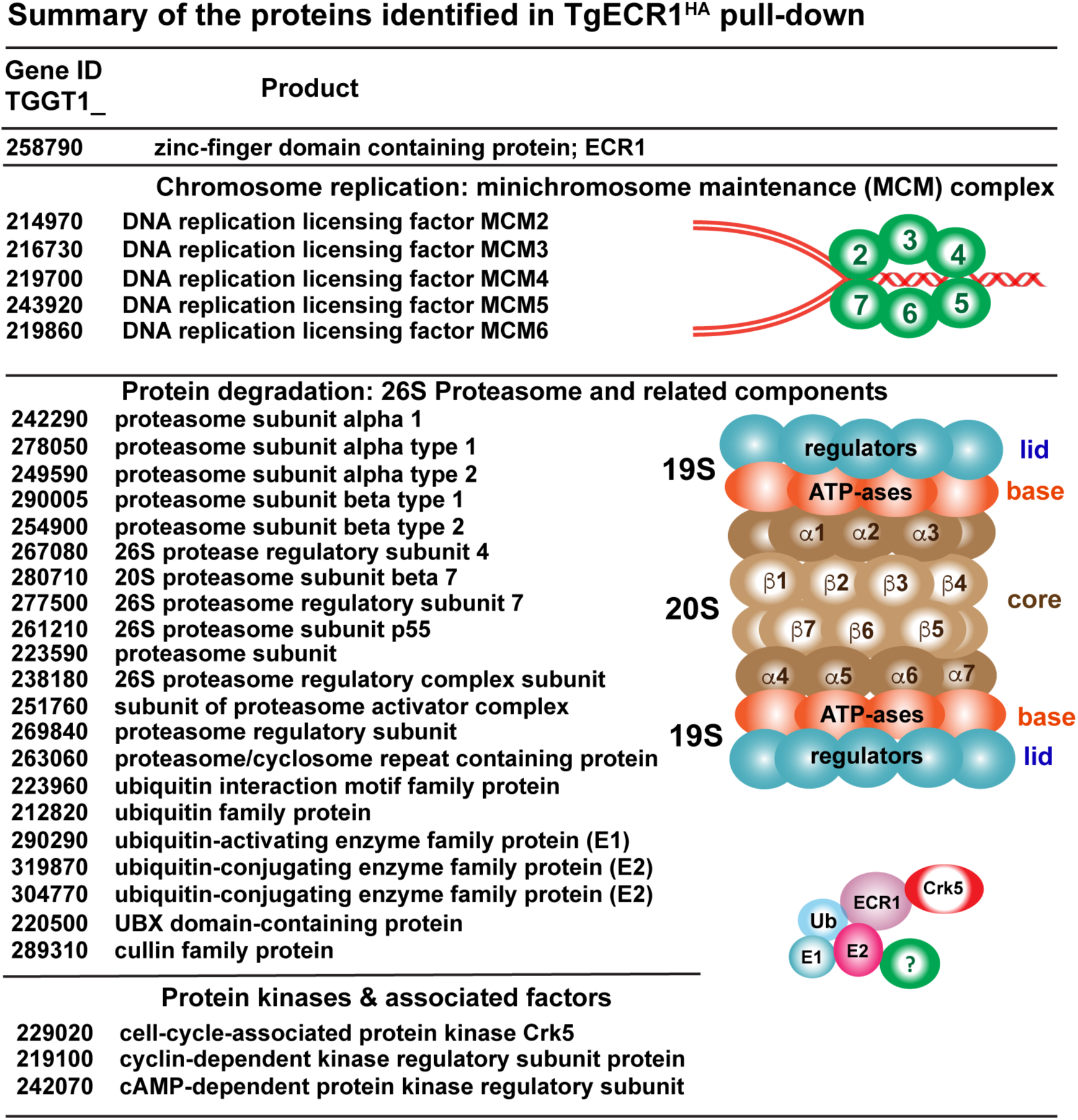
ECR1 proteomics. Proteomic analysis of proteins pulled down with ECR1^HA^ incudes five minichromosome maintenance complex proteins (MCM2 to MCM6) required for DNA replication. The ECR1 proteome also contained a number 20S core alpha- and beta-subunits, 19S regulatory particle proteins, as well as members of ubiquitination machinery, e.g. ubiquitin-activating enzyme, two ubiquitin-conjugating enzymes. ECR1 also interacted with protein kinases and associated factors including a CDK-related kinase, TgCrk5 (16).

CDK protein kinases are key factors in cell cycle checkpoint control (17), therefore, we further investigated TgCrk5 expression in the tachyzoite cell cycle. The TgCrk5 protein was C-terminally epitope tagged with 3xHA by genetic knock-in and IFA analysis of TgCrk5^HA^ revealed a similar cell cycle profile as ECR1 with maximum protein expression in S phase and low to near undetectable levels in G1 parasites (Fig. 6A, also see mRNA profile Fig. 7A). Like ECR1, TgCrk5^HA^ is nuclear factor that concentrates in the centrocone where it co-localized with tsECR1^Myc^ in a dual-epitope tagged strain generated by serial knock-in (Fig. 6B, tsECR1^Myc^/TgCrk5^HA^, 34^°^C images). Utilizing the dual-tagged strain, tsECR1^Myc^ interaction with TgCrk5^HA^ at the permissive temperature (34^°^C) was confirmed by co-immunoprecipitation (co-IP) of whole cell lysates (Fig. 6C). Finally, the expression of TgCrk5^HA^ in the dual-tagged strain at 40^°^C was examined revealing TgCrk5^HA^ protein levels largely remained associated with parasite chromosome material at 40^°^C (Fig. 6D), although TgCrk5^HA^ centrocone staining was abolished consistent with the disruption of this structure at high temperature. Staining with TgPCNA1 antibodies distinguishes nuclear chromatin from aggregated plastid DNA in these defective parasites (images 16, 17), and as expected TgCrk5 co-localizes with TgPCNA1 in the tsECR1 parasites (data not shown).

**FIG. 6.**
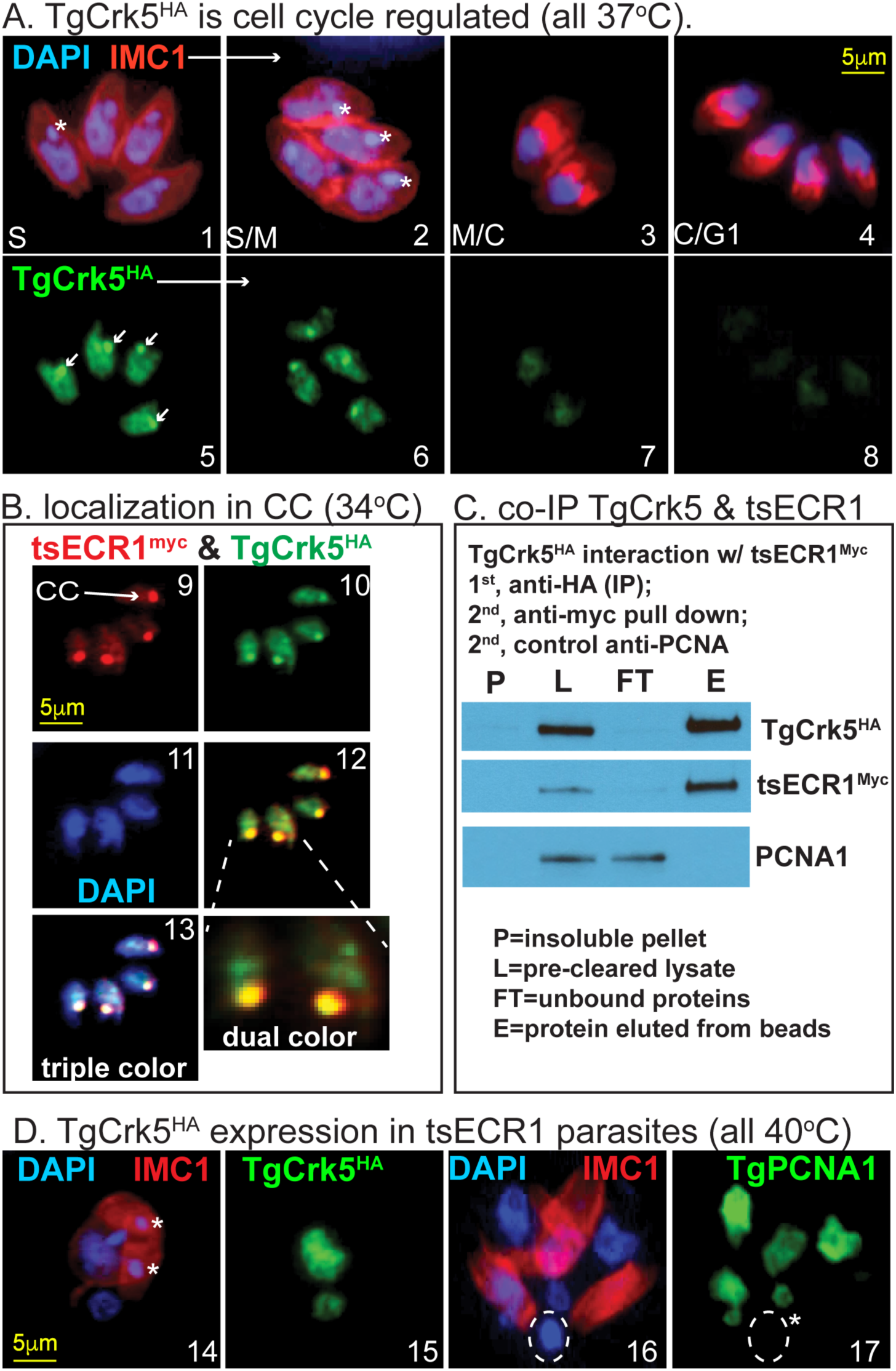
TgCrk5 is associated with ECR1 during tachyzoite replication. **[A].** TgCrk5^HA^ transgenic parasites were co-stained with aHA, aIMC1 and DAPI and representative images of parasites in S through early G1 phase of the next cell cycle are shown. Images 1-4, merge of αIMC1 and DAPI staining showing the mother IMC and daughter buds in relation to genomic and plastid DNA. Note the daughter buds formed in image 3 precede nuclear division, whereas, nuclei in the four parasites in image 4 are now packaged into the nearly mature daughters representative of late cytokinesis and early G1 cell cycle time periods. Stars indicate plastid DNA. Images 5-8, co-staining of the parasites in images 1-4 with αHA revealing the cell cycle expression patterns of TgCrk5^HA^. The maximum expression in S phase and low to near undetectable expression in late cytokinetic/early G1 parasites are consistent with the encoded mRNA cell cycle profile (Fig. 7A). Localization of TgCrk5^HA^ in the centrocone is indicated by arrows in image 5. [B.] TgCrk5 was epitope tagged with 3xHA in tsECR1^Myc^ transgenic parasites. IFA analysis (images 9-13) of this new dual epitope tagged strain demonstrated co-localization of tsECR1^Myc^ and TgCrk5^HA^ in the parasite centrocone (CC) and nucleus at 34^°^C. Note the expression level of TgCrk5^HA^ was higher than tsECR1^Myc^ in these parasites. Image 12=merged staining results for TgCrk5^HA^ and tsECR1^Myc^ expression with higher magnification the two centrocone structures indicated. Image 13=merged staining results for aHA, aMyc, and DAPI stain used to indicate nuclear DNA. **[C.]** Immunoprecipitation of TgCrk5^HA^ confirms interaction with tsECR1^Myc^ in parasites grown at 34^°^C. Western analysis reveals that virtually all cellular tsECR1^Myc^ protein was pulled down with TgCrk5^HA^, while the abundant nuclear factor TgPCNA1 was not pulled down. **[D.]** IFA analysis of the dual tagged TgCrk5^HA^/tsECR1^Myc^ strain at 40^°^C. Similar to [A.] parasites were co-stained for DNA and IMC1 (image 14 and 16) and then for TgCrk5^HA^ (image 15) or TgPCNA1 (image 17). Stars in images 14 and 17 indicate plastid DNA, which in images 16 and 17 have aggregated into a single deposit (dashed circle) that does not contain TgPCNA1. Magnification scale bars for each image series **[A. B. and C.] a**re indicated once.

### Loss of tsECR1 affects the stability of TgMCM2

In *T. gondii* tachyzoites, mRNAs encoding the MCM_2-7_ proteins are cyclically expressed and tightly coordinated with maximum expression in late G1 phase (Fig. 7A). By contrast, ECR1 and TgCrk5 mRNAs and protein levels reached maximum expression later in S phase (Figs. 3 & 4 and 7A). To confirm the periodic expression of representative MCM factors in *T. gondii* at the protein level, we epitope tagged TgMCM2 with 3xHA by genetic knock-in and evaluated cell cycle expression of this protein in randomly growing tachyzoites (Fig. 7B). Roughly half of transgenic tachyzoites in a randomly growing population expressed TgMCM2^HA^ with peak expression during the transition from G1 into S phase (Fig. 7B) consistent with the encoded mRNA (Fig. 7A). During daughter bud formation TgMCM2^HA^ protein decreased below IFA detection (Fig. 7B). We then investigated the expression of TgMCM2^HA^ in parasites carrying the tsECR1 mutation (Fig. 7C,D). In these ts-mutant parasites, TgMCM2^HA^ was detected at high levels in more than 80% of the population (see Fig. 7C graph) and was colocalized to the masses of chromosome DNA that accumulated in the mother parasite at 40^°^C (Fig. 7B, bottom panel). The increase in TgMCM2^HA^ levels at 40^°^C in the tsECR1 parasites was in part due to greater TgMCM2^HA^ stability in tsECR1 parasites (which lack ECR1 at high temperature) as compared to parasites that express native ECR1 (Fig. 7D). Protein stabilization was not a generalized phenotype of the loss of tsECR1 or a consequence of high temperature as there was no increased stabilization of the TgCrk5 partner in tsECR1 parasites grown at 40^°^C (Fig. 7D) nor did we detect changes in the stability of αtubulin. It should be noted that we did not detect stable association of MCM2^HA^ with tsECR1^Myc^ at permissive temperatures using co-IP methods that successfully confirmed tsECR1^Myc^/TgCrk5^HA^ interaction in these parasites (results not shown). If MCM2 is an ECR1 substrate, then the interaction is expected to be transient and likely only detectable using substrate trapping techniques (18).

**FIG. 7.**
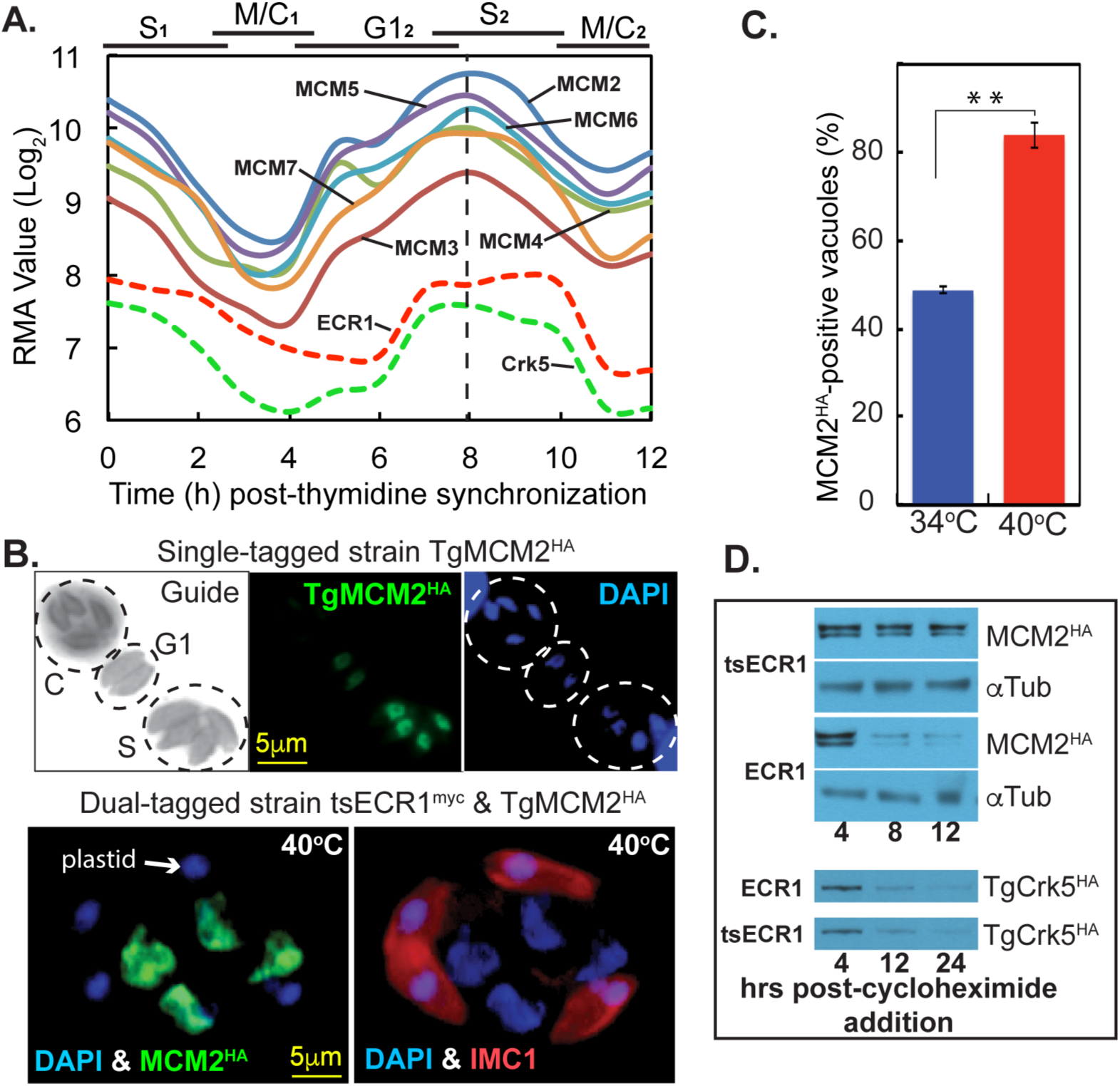
Disruption of ECR1 stabilizes TgMCM2 expression. **[A.]** TgMCM_2_-7 mRNAs levels in synchronized tachyzoites (21) revealed nearly identical cyclical profiles (dashed vertical line indicates shared peak expression). Cyclical profiles for ECR1 and TgCrk5 mRNAs are included to show relative cell cycle timing; TgMCM_2-7_ mRNAs peak earlier that ECR1 and TgCrk5 mRNAs. **[B.]** Cell cycle analysis of TgMCM2^HA^ expression: co-stains aHA (TgMCM2^HA^, green) and DAPI (DNA, blue). Guide panel is a non-colorized/inverted IMC1 expression showing three vacuoles in different cell cycle phases as indicated (C=budding vacuole, and late G1 and S phase vacuoles). TgMCM2^HA^ maximum and minimum expression occurred in S phase and mitosis, respectively, which was consistent with the encoded mRNA cell cycle profiles in [A.]. Magnification scale bar is indicated. **[C.]** IFA analysis of TgMCM2^HA^ tagged in a tsECR1^Myc^ strain at 40^°^C. Note, masses of nuclear DNA (DAPI, blue) in the parasite are associated with high levels of TgMCM2^HA^ (green), whereas abnormal buds (IMC1, red) only contain plastid DNA (DAPI, blue). Parasites positive for TgMCM2^HA^ in 100 randomly selected vacuoles nearly doubled when the tsECR1^Myc^ strain was shifted to 40^°^C for 24 h. At 34^°^C, the ∼50% of parasites expressing TgMCM2^HA^ correspond to late G1 through S phase as shown in [C.]. **[D.]** Two dual epitope-tagged tachyzoite strains (tsECR1^Myc^ with TgMCM2^HA^ or with TgCrk5^HA^) or the single TgMCM2^HA^ strain were shifted to 40^°^C, treated with 200 μM cycloheximide to block total protein synthesis, and Western analysis of TgMCM2^HA^ and TgCrk5^HA^ protein levels determined at the times indicated. Tubulin is naturally stable and included here to demonstrate equal loading. Note that in tsECR1^myc^ parasites at 40^°^C, TgMCM2^HA^ protein stability was increased, while the stability of TgCrk5^HA^ was unaffected.

## DISCUSSION

Compared to yeast and mammalian somatic cell proliferation, cell division of apicomplexan asexual life cycle stages is unique in a number of ways. Progeny production from a single mother cell varies widely and is a characteristic of each species and stage, which indicates parasite genetics is responsible for these differences. The ability to suspend cytokinesis (and nuclear division in some cases, e.g. *Sarcocystis neurona*)(19), while chromosomes and nuclei are reduplicated is remarkable as is the cytoskeletal complexity that must be assembled to produce infectious daughter parasites. The knowledge that Apicomplexa replication is unusual was an early discovery of morphogenic studies (e.g. ref. (4) and we have expected the molecular control of apicomplexan cell division to also have distinct features. Through comparative genomics and experimental genetics major checkpoint mechanisms of the *T. gondii* tachyzoite stage were recently defined (16). For a single cell eukaryote, there are a large number of Crk protein kinases required for tachyzoite replication, including the ECR1/TgCrk5 mechanism described here (see Fig. 8 model)(16). A single TgCrk controls G1 progression, while up to four TgCrks regulate karyokinesis and/or cytokinesis in the tachyzoite (16). In contrast to a recent suggestion (20), there are no canonical cyclins (A-E type) in the major Apicomplexa families; cyclins in these parasites are atypical as are many of the TgCrks (16). Oscillating cyclins program checkpoint kinases in higher eukaryotes, however, experimental evidence indicates some TgCrks lack cyclin partners. The discovery here that TgCrk5 is periodically expressed in the tachyzoite is unusual for a potential checkpoint kinase, and dynamic expression extends to mitotic kinases TgCrk4 and TgCrk6 (16). Perhaps because they are periodic, TgCrk4 and TgCrk6 lack detectable cyclin partners, and it is possible that the cyclically expressed TgCrk5 will also lack a cyclin partner.

**FIG. 8.**
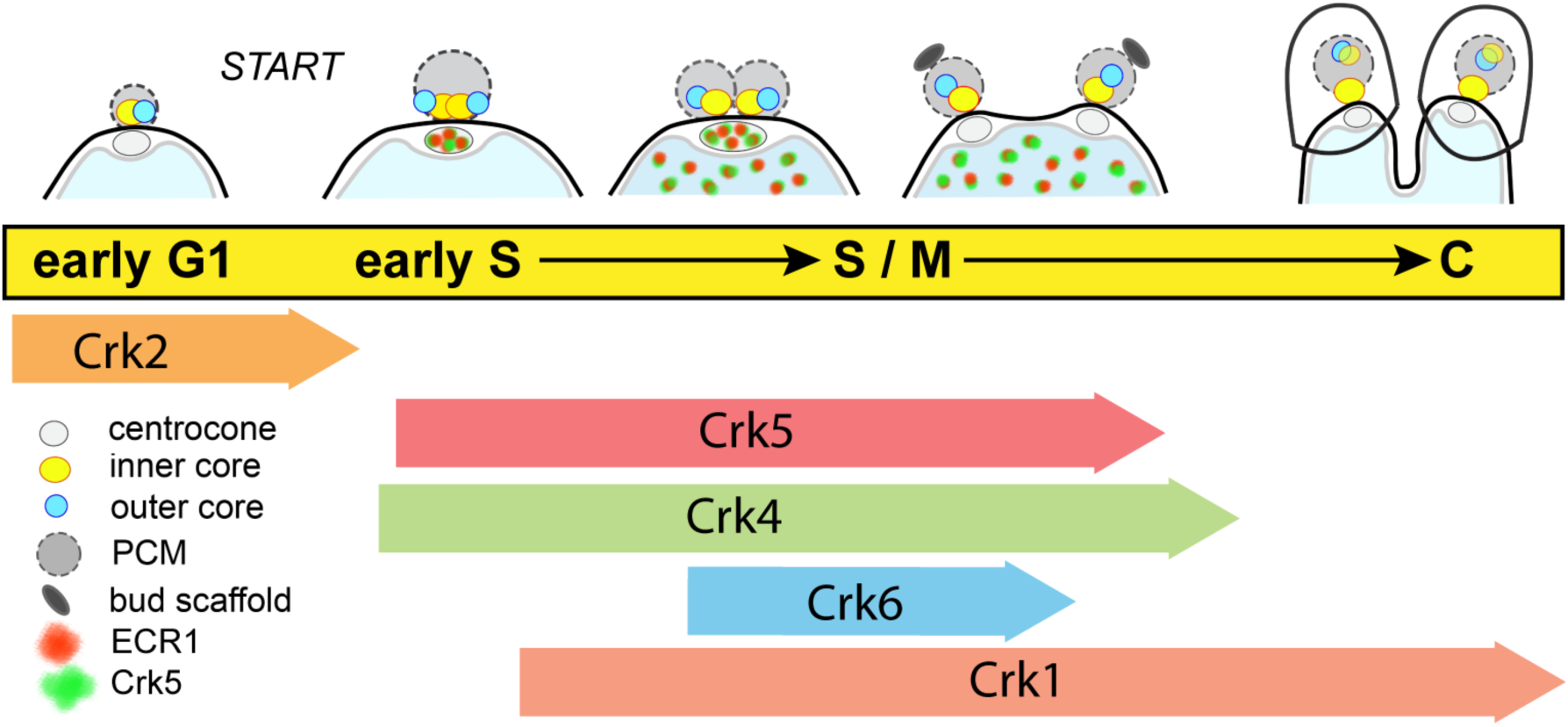
Major control mechanisms of the *T. gondii* tachyzoite cell cycle. Major cell cycle transitions are indicated by conventional phase designations overlaid by the sequential timing of centrosome outer and inner cores and centrocone duplication in dividing tachyzoites. PCM=Pericentrosomal matrix surrounding the centrosome cores. The cell cycle periods corresponding to the mitotic events diagramed above are linked to the timing of proposed functions of TgCrks indicated below (16). Localization and expression timing of ECR1 (green) and TgCrk5 (red) that begins in the centrocone and shuttles to the nucleus is indicated. Note that four TgCrks regulating S phase and mitosis progression is unusual and distinct from well-studied eukaryotic models (yeast and human cells) that require 1-2 CDKs for these same processes (16).

The enzymology of DNA synthesis of eukaryotes is well established and conserved (15). The DNA helix is unwound by an active helicase assembled from six core MCM_2-7_ proteins and new DNA strands are then produced by DNA polymerases supported by accessory factors (see Fig. S3B model). Apicomplexa parasites possess most of this DNA synthetic machinery (Fig. S3A summary), which includes a requirement for a topoisomerase as we have demonstrated here (Fig. 1). However, more than the basic machinery is conserved as the topology of G1 through S phase progression in these parasites has parallels to higher eukaryotes. Like other eukaryotes, ordered gene expression (protein synthesis first, DNA replication factors second) in the apicomplexan G1 phase (21, 22) prepares the way for DNA synthesis. The ability to synchronize *T. gondii* tachyzoites at the G1/S boundary (6) indicates there is a START-like checkpoint (likely controlled by TgCrk2) (16) controlling commitment to DNA synthesis and the faithful reduplication of nuclei in apicomplexans indicate some type of “copy once-only once” rule is also active (Fig. 8). In human cells, the E2F/CDK4,6/retinoblastoma controls the G1 pathway (23) and multiple protein regulators beyond the CDK/cyclin mechanisms coordinate each step of chromosome replication (15). Orthologs of these human cell cycle regulators are not present in Apicomplexa genomes (Fig. S3A), although there is precedence for lineage-specific proteins to produce similar cell cycle topologies (24). Therefore, the question is not if but what are the cell cycle regulators in the Apicomplexa.

The ubiquitin-proteasome system serves a critical role in the regulation of chromosome replication, and here the basic regulatory factors are conserved and active in the Apicomplexa (25). A full complement of ubiquitin-conjugating factors and proteasome components are present and a recent global survey of ubiquitin-conjugated proteins in *T. gondii* tachyzoites (25) determined that ∼35% of proteins with this modification are cell cycle regulated (21). Relevant to this study is a remarkable peak in the ubiquitination and phosphorylation of proteins in late S phase of the tachyzoite cell cycle (25). The RING/TRAF zinc finger factor, ECR1, characterized here may be part of this system (Fig. 8 and Fig. S3B). ECR1 interaction with many components of the proteasome as well as two E2 conjugating enzymes, indicates this factor could be a unique E3 ligase, which will require future biochemical experiments to confirm. Furthermore, ECR1 potential interaction with MCMs along with the stabilization of MCM2 when ECR1 is depleted (Fig. 7) suggests ECR1 may specifically function in the licensing of chromosome origins of replication in the tachyzoite. The timing of ECR1 expression in S phase and early mitosis, which is after the increase in MCM_2-7_ expression in G1 (Fig. 7A), suggests this factor is not responsible for the initial licensing of origins. However, all eukaryotes need a mechanism that prevents re-licensing during active chromosome replication. In human cells, the protein geminin serves this function, and shares a similar biology to ECR1 described here. Geminin is only active in S phase through early mitosis and interacts with and is regulated by a CDK-related kinase (human CDK2) through a reversible ubiquitination mechanism (26-28). Like ECR1, geminin has functions beyond regulating chromosome replication that involve ensuring proper centrosome duplication and spindle formation (29). It remains a possibility that apicomplexan parasites will have a divergent geminin-like protein and the ECR1 proteome (Dataset S1) includes 24 unknown proteins with 9 proteins encoded by an mRNA that has maximum expression in S-phase (gene records for all ECR1 interactors can be obtained at ToxoDB). These 9 proteins also possess coiled-coil domains, which is the key structural feature of human geminin.

The presence of multiple microtubule organizing centers in apicomplexan parasites (MTOCs; centrosome/centrocone and apical complex)(30, 31) sets these parasites apart from well studied eukaryotic models (yeast and mammalian cells) that have a single MTOC during interphase (32). Our recent discovery of a novel bipartite centrosome with independent core centers that separately regulate cytokinesis and karyokinesis adds further physical and molecular complexity to apicomplexan cell division (2). The centrocone and apical complex MTOCs are conserved across Apicomplexa lineages with few if any parallels in better studied eukaryotes. Contained within the nuclear membrane, the enigmatic centrocone (33) is present throughout apicomplexan cell division (34) and in *T. gondii* tachyzoites this structure duplicates and then separates starting in late S phase (Fig. 8)(2). The appearance of the ECR1/TgCrk5 complex in the centrocone prior to duplication suggests these factors may have a role in preparing this structure for mitosis. The mitotic spindle originates in the centrocone (35) and through this structure nuclear and cytoplasmic mitotic processes are also physically connected (2, 14). Our and other studies show that this involves the centrosome inner core on the cytoplasmic face (2) and the kinetochore on the nuclear side (14). In the tachyzoite metaphase these mitotic structures are aligned in a typical linear array with the centrocones anchoring each pole (2). There is clear evidence from studies in other eukaryotes that spindle structures house cell cycle regulatory proteins. Classically, the transitory association of CDK1 and mitotic cyclin complex in the yeast spindle pole body is associated with activation of this checkpoint mechanism (36, 37). In budding and fission yeast regulatory protein association with the spindle pole body is also required for mechanisms of mitotic exit, which includes factors of the MEN and SIN complexes (38, 39). Until our study here only one protein, MORN1, had been localized within the apicomplexan centrocone, which may be functionally, if not structurally equivalent to the yeast spindle pole body. The MORN1 marker has permitted the morphogenesis of this important structure to be defined in several apicomplexan parasites (34). We have established here that ECR1 and TgCrk5 are new markers of the centrocone and their presence in the centrocone demonstrates that potential checkpoint controls are housed in this unique spindle organizing structure. The importance of the ECR1/TgCrk5 complex for the duplication of the centrocone and maintaining the physical connection between nuclear and cytoplasmic mitotic centers was revealed in the lethal arrest of the tsECR1 ts-mutant (Fig. 4). At high temperature, inner and outer centrosome cores, the centrocone, and the kinetochores became uncoupled and all three structures showed reduced or abnormal duplication (Fig. 4). The stability of TgCrk5 is unaffected by the loss of ECR1, however, localization of TgCrk5 in the centrocone requires this factor. Altogether, these results indicate that the localization of ECR1/TgCrk5 in the centrocone and nucleus likely does not represent inactive versus active forms and instead the results suggest the ECR1/TgCrk5 complex has multiple temporally regulated functions vital for tachyzoite cell division. The association of this complex prior to centrocone duplication and spindle formation may be required to prepare this structure for mitosis and to establish communication between the newly duplicated centrosome and nuclear kinetochore, while nuclear localization of ECR1/TgCrk5 could be required to prevent re-licensing of origins during chromosome replication. There is evidence from phosphoproteomics that ECR1 is phosphorylated as is TgCrk5 (see ToxoDB gene records) opening the possibility that ECR1 function may require TgCrk5 phosphorylation. Deciphering the full molecular details of the ECR1/TgCrk5 mechanisms will require further molecular dissection as well as the understanding what other proteins participate including the identity of substrates for both the potential ubiquitination and phosphorylation activities. In addition, ECR1 and TgCrk5 provide new tools to probe the scope of critical cell cycle functions housed in the centrocone compartment. While there is more to be done, what this study, and the related investigation of *T. gondii* checkpoints (16) shows, is that our understanding of the molecular basis of apicomplexan cell division is beginning to finally catch up to the remarkable morphogenic processes discovered decades ago.

## MATERIALS AND METHODS

### Parasite cell culture and flow cytometry

Parasites were grown in human foreskin fibroblasts (HFF) as described (40). The ts-mutants 11-51A1 and 13-64D5 were obtained by chemical mutagenesis (7) of the *RHΔhxgprt* parasite strain (41). All transgenic lines used in this study were produced in the *RHΔku80Δhxgprt* strain (42); see Dataset S2 for full list of genes, primers and transgenic strains. Growth measurements were performed using parasites pre-synchronized by limited invasion as previously described (10, 43). Parasite vacuoles in the infected cultures were evaluated over various time periods with average vacuole sizes determined at each time point from 50-100 randomly selected vacuoles.

Parasite nuclear DNA content was determined by flow cytometry using propidium iodide (PI) (Sigma, St. Louis, MO) as described previously (6). Briefly, filter-purified tachyzoites were fixed in 70% (v/v) ethanol, pelleted at 3000 x g, and then suspended in phosphate buffered saline (6 × 10^6^ parasites/ml) and stained with PI (0.2 mg/ml final concentration in 0.5 ml total volume). RNase cocktail (250U; combination of RNase A, RNase T1, Ambicon Corp., Austin, TX) was added and the parasites incubated in the dark at room temperature for 30 min. Nuclear DNA content was measured based on PI (FL-2) fluorescence using a FACS-Calibur flow cytometer (Becton-Dickinson Inc., San Jose, CA). Fluorescence was collected in linear mode (10,000 events) and the results were quantified using CELLQuest^™^ v3.0 (Becton-Dickinson Inc.).

### Immunofluorescence assays and Western analysis

Confluent HFF cultures on glass coverslips were infected with parasites for the indicated times. Infected monolayers were fixed, permeabilized and incubated with antibody as previously described (10). The following primary antibodies were used; αTgCenH3 rabbit polyclonal (kinetochore stain) (14), αHA rat monoclonal (3F10, Roche Applied Sciences), αMyc rabbit monoclonal (71D10, Cell Signaling Technology), αHuman Centrin 2 rabbit polyclonal (centrosome outer core stain) (11), αMORN1 rabbit polyclonal (centrocone and basal complex stains, kindly provided by Dr. Marc-Jan Gubbels, Boston College, MA), αIMC1 mouse monoclonal or rabbit polyclonal (parasite shape and internal daughter bud stains, kindly provided by Dr. Gary Ward, University of Vermont, VT). All Alexa-conjugated secondary antibodies (Molecular Probes, Life Technologies) were used at dilution 1:500. Coverslips were mounted with Aquamount (Thermo Scientific), and viewed on Zeiss Axiovert Microscope equipped with 100x objective.

For Western analysis, purified parasites were washed in cold PBS and collected by centrifugation. Total lysates were obtained by suspending the parasite pellets with Laemmli loading dye, heated at 75°C for 10 min, and briefly sonicated. After separation on the SDS-PAGE gels, proteins were transferred onto nitrocellulose membrane and probed with the following antibodies: αHA rat monoclonal (3F10, Roche Applied Sciences), αMyc rabbit monoclonal (Cell Signaling Technology) and αTubulin mouse polyclonal (12G10, kindly provided by Dr. Jacek Gaertig, University of Georgia). After incubation with secondary HRP-conjugated antibodies against amouse, arabbit or αrat primary antibodies, proteins were visualized using the Western Lightning^®^ Plus-ECL chemiluminescence reagents (PerkinElmer).

### Identification of temperature sensitive mutations

The ts-mutant 11-51A1 was complemented using ToxoSuperCos cosmid library as published (7, 10, 11). Briefly, ts-mutant parasites were transfected with cosmid library DNA (50 μg DNA/5 × 10^7^ parasites/transfection) in twenty independent electroporations. After two consecutive selections at 40°C, parasites were selected by the combination of high temperature and 1 μM pyrimethamine and then genomic DNA was isolated for marker-rescue (7). The rescued genomic inserts were sequenced in order to map the rescue locus to the *T. gondii* genome (ToxoDB). To resolve the contribution of individual genes in the recovered locus, we transformed the ts-mutant 11-51A1 with DNA fragments representing four genes included in the locus: TGME49_258810 (gene 2), TGME49_258800 (gene 3), TGME49_258790 (gene 4) and TGME49_258780 (gene 5) (Fig 2A). The gene-specific DNA fragments were PCR amplified from genomic DNA of the parental strain *RHΔhxgprt* (primers used are listed in the Dataset S2) and transfected into 1x10^7^ parasites using 6-10 μg of purified DNA. Transfected 11-51A1 ts-mutant populations were grown at permissive temperature overnight and the selected at high temperature. Successful genetic rescue was only achieved with amplified genomics DNA spanning the TGME49_258790 gene. The defective gene in ts-mutant 13-64D5, which as a similar phenotype to ts-mutant 11-51A1, was identified by whole-genome sequencing followed by selected fosmid (carrying genomic inserts) complementation using a published strategy (12). Whole genome DNA libraries for ts-mutant 13-64D5 were prepared, sequenced, and analyzed for single nucleotide variation according to published methods (12); sequencing of ts-mutant 13-64D5 resulted in >150x genome coverage and identified 16 non-synonymous mutations. Based on the sequencing results, a stepwise fosmid complementation strategy was developed for ts-mutant 13-64D5 (Fig. 2B), which revealed the defective gene in 13-64D5 was a *T. gondii* ortholog (TGME49_312230) of eukaryotic DNA topoisomerase II.

### Generation of transgenic tachyzoite strains

#### Endogenous tagging by genetic knock-in technique

Selected *T. gondii* proteins were tagged with a triple copy of the HA or Myc tag by previously described genetic knock-in protocols (42). PCR DNA fragments encompassing the 3’-end of the gene of interest (GOI) were used to construct the plasmids pLIC-GOI-HA_3X_/dhfr-DHFR-TS, pLIC-GOI-HA_3X_/dhfr-HXGPRT or pLIC-GOI-Myc_3X_/dhfr-DHFR-TS and the constructs were transfected into *RHΔku80Δhxprt* strain deficient in non-homologous recombination and the HXGPRT purine salvage protein (42). The double-tagged transgenic lines were established by sequential selection under alternative drug selection with cloning and dual epitope tag expression was verified by IFA. To reconstruct the tsECR1 phenotype in a clean genetic background, a strain expressing the ts-ECR1^Myc^ was generated using the same C-terminal knock-in strategy (Fig. S1B). The DNA insert used to build the tsECR1 knock-in plasmid was chemically synthesized (GenScript USA Inc, Piscataway, NJ) to incorporate the L413H mutation along with a 3xMyc epitope tag while also modifying the amino acid coding to prevent recombination events by-passing the ts-mutation. Phenotypic regeneration of the new tsECR1 strain was verified by IFA and Western analysis (Fig. S2).

#### Ectopic expression of epitope-tagged ECR1

The coding sequence of ECR1 was amplified from parental *RHΔhxgprt* or tsECR1 cDNA libraries (see Dataset S2 for primers design) and the DNA-fragments were cloned into the pDEST_gra-Myc_3X_/sag-HXGPRT vector by recombination (Gateway cloning, Life Technologies), with the 3xMyc tag fused to the C-terminus. Constructs were transformed into parental *RHΔhxgprt* strain and selected with mycophenolic acid and xanthine. The resulting ECR1 and tsECR1 clones were evaluated by Western analysis (Fig. S1A).

### Phylogenetic analysis of ECR1 and TgTopoII

Amino acid sequence of ECR1 was compared to all non-redundant CDS translations in ncbi.org and in Eupath.org and we found strong ECR1 orthologs in coccidian branch of Apicomplexa and divergent orthologs in chromerids and filamentous fungi. Convincing TgTopo-II orthologues where identified in all major eukaryotic groups by pBLAST (ncbi.org). Evolutionary analyses were conducted in Phylogeny.fr (44). The bootstrap consensus tree inferred from 100 replicates represents the evolutionary history of ECR1 and TgTopo-II families analyzed by Advanced method based on WAG substitution model (44). To build phylogenetic tree of ECR1-related factors, human PDZRN3 protein containing RING and Sina domains was used as an outgroup.

### Proteomics and validation by co-immunoprecipitation

The ECR1 protein fused to a 3xHA tag on C-terminus was used for proteomic analysis. ECR1^HA^ expressing tachyzoites were grown at MOI=2 for 26 h at 37^°^C and 5.48 × 10^9^ parasites were collected before the lysis to ensure a sufficient sample of S phase parasites. To prepare the parasite lysate, the parasite pellet was incubated 60 min at 4°C in lysis buffer (0.1% [v/v] Nonidet P-40, 10 mM HEPES pH 7.4, 150 mM KCl, plus protease inhibitors and phosphatase inhibitors) with rotation, and subjected to six freeze-thaw cycles (snap freezing in liquid nitrogen bath and thawing ice-water bath) with vortex for 1 min at 4°C before freezing. The parasite lysate was centrifuged 12,000×g for 30 minutes at 4°C, and the clarified lysate was used for co-immunoprecipitation.ECR1^HA^ was immunoprecipitated using mouse monoclonal anti-HA-tag magnetic beads (μMACS Anti-HA Microbeads; Miltenyi Biotec). The parasite lysate was incubated with anti-HA magnetic beads overnight at 4°C with rotation. After the beads were washed 4 times with cold wash buffer 1 (150 mM NaCl, 1% NP-40, 0.5% sodium deoxycholate, 0.1% SDS, 50 mM Tris-HCl pH 8.0) and once with wash buffer 2 (20 mM Tris-HCl pH 7.5) by using μ Column (prewashed with buffer containing 0.1% [v/v] Nonidet P-40, 10 mM HEPES pH 7.4, 150 mM KCl) in the magnetic field of the separator, the bound proteins (co-IP complexes) were eluted from the magnetic beads by applying 50 μl of pre-heated 95 °C hot Laemmli’s sample buffer to the column, then were separated by SDS-PAGE (Mini-PROTAN^®^ TGX 4-20%; BioRad), and stained with Coomassie blue (GelCode Blue Stain Reagent; Pierce). Each sample lane was cut into 24 slices and separately analyzed by LC-MS/MS as previously described (8). Briefly, proteins reduced and alkylated with TCEP and iodoacetamide were digested with trypsin, and run sequentially on Acclaim PepMap C18 Nanotrap column and PepMapRSLC C18 column (Dionex Corp). Raw LC-MS/MS data was collected using Proteome Discoverer 1.2 (Thermo Scientific) and proteins were searched in Toxo_Human Combined database using in-house Mascot Protein Search engine (Matrix Science) and the final list was generated in the Scaffold 5.5.1 (Proteome Software) with following filters: 99% protein probability with a minimum 2 peptides with >95% peptide probability.

Co-IP of tsECR1^Myc^ and TgCrk5^HA^ was performed as following. Dual tagged TgRHΔku80-tsECR1^Myc^::TgCrk5^HA^ parasites were grown at 34^°^C for 40 h and 2×10^8^ parasites were collected, lysed and immunoprecipitated on anti-HA beads (Medical and Biological Laboratories, Japan) as described previously (2). Protein extracts were rotated with beads at RT for 1 h, washed beads were heated for 10 min at 75^°^C in Laemmli sample buffer to elute bound proteins. Proteins were separated on Mini-PROTAN^®^ TGX 4-20% gels (BioRad), transferred to nitrocellulose membrane, and incubated with αMyc and αHA antibodies. After incubation with secondary HRP-conjugated antibodies (Jackson ImmunoResearch Lab, West Grove, PA), proteins were visualized using Western Lightning^®^ Plus-ECL chemiluminescence reagents (PerkinElmer).

## ACKNOWLEDGEMENTS

*T. gondii* genomic and/or cDNA sequence data were accessed via http://ToxoDB.org. The authors would like to thank Dr. Sumiti Vinayak (University of Georgia-Athens, GA) for the help with whole genome sequence analysis.

## REFERENCES

1. Bhagavathula AS, Elnour AA, Shehab A. 2016. Alternatives to currently used antimalarial drugs: in search of a magic bullet. Infect Dis Poverty 5: 103.

2. Suvorova ES, Francia M, Striepen B, White MW. 2015. A novel bipartite centrosome coordinates the apicomplexan cell cycle. PLoS Biol 13: e1002093.

3. Gubbels MJ, White M, Szatanek T. 2008. The cell cycle and Toxoplasma gondii cell division: tightly knit or loosely stitched? Int J Parasitol 38: 1343–58.

4. Sheffield HG, Melton ML. 1968. The fine structure and reproduction of Toxoplasma gondii. J Parasitol 54: 209–26.

5. Radke JR, Striepen B, Guerini MN, Jerome ME, Roos DS, White MW. 2001. Defining the cell cycle for the tachyzoite stage of Toxoplasma gondii. Mol Biochem Parasitol 115: 165–75.

6. Radke JR, White MW. 1998. A cell cycle model for the tachyzoite of Toxoplasma gondii using the Herpes simplex virus thymidine kinase. Mol Biochem Parasitol 94: 237–47.

7. Gubbels MJ, Lehmann M, Muthalagi M, Jerome ME, Brooks CF, Szatanek T, Flynn J, Parrot B, Radke J, Striepen B, White MW. 2008. Forward genetic analysis of the apicomplexan cell division cycle in Toxoplasma gondii. PLoS Pathog 4: e36.

8. Suvorova ES, Croken M, Kratzer S, Ting LM, de Felipe MC, Balu B, Markillie ML, Weiss LM, Kim K, White MW. 2013. Discovery of a splicing regulator required for cell cycle progression. PLoS Genet 9: e1003305.

9. Radke JR, Guerini MN, White MW. 2000. Toxoplasma gondii: characterization of temperature-sensitive tachyzoite cell cycle mutants. Experimental parasitology 96: 168–77.

10. Suvorova ES, Lehmann MM, Kratzer S, White MW. 2012. Nuclear actin-related protein is required for chromosome segregation in Toxoplasma gondii. Molecular and biochemical parasitology 181: 7–16.

11. Suvorova ES, Radke JB, Ting LM, Vinayak S, Alvarez CA, Kratzer S, Kim K, Striepen B, White MW. 2013. A nucleolar AAA-NTPase is required for parasite division. Mol Microbiol 90: 338–55.

12. Vinayak S, Brooks CF, Naumov A, Suvorova ES, White MW, Striepen B. 2014. Genetic manipulation of the Toxoplasma gondii genome by fosmid recombineering. MBio 5: e02021.

13. Matsuzawa SI, Reed JC. 2001. Siah-1, SIP, and Ebi collaborate in a novel pathway for beta-catenin degradation linked to p53 responses. Mol Cell 7: 915–26.

14. Brooks CF, Francia ME, Gissot M, Croken MM, Kim K, Striepen B. 2011. Toxoplasma gondii sequesters centromeres to a specific nuclear region throughout the cell cycle. Proceedings of the National Academy of Sciences of the United States of America 108: 3767–72.

15. Sacco E, Hasan MM, Alberghina L, Vanoni M. 2012. Comparative analysis of the molecular mechanisms controlling the initiation of chromosomal DNA replication in yeast and in mammalian cells. Biotechnol Adv 30: 73–98.

16. Alvarez C, Suvorova ES. 2017. A checkpoint roadmap for the complex cell division of Apicomplexa parasites. bioRxiv https://doiorg/101101/104646.

17. Bartova I, Koca J, Otyepka M. 2008. Functional flexibility of human cyclin-dependent kinase-2 and its evolutionary conservation. Protein Sci 17: 22–33.

18. Kim TY, Siesser PF, Rossman KL, Goldfarb D, Mackinnon K, Yan F, Yi X, MacCoss MJ, Moon RT, Der CJ, Major MB. 2015. Substrate trapping proteomics reveals targets of the betaTrCP2/FBXW11 ubiquitin ligase. Mol Cell Biol 35: 167–81.

19. Vaishnava S, Morrison DP, Gaji RY, Murray JM, Entzeroth R, Howe DK, Striepen B. 2005. Plastid segregation and cell division in the apicomplexan parasite Sarcocystis neurona. J Cell Sci 118: 3397–407.

20. Francia ME, Striepen B. 2014. Cell division in apicomplexan parasites. Nat Rev Microbiol 12: 125–36.

21. Behnke MS, Wootton JC, Lehmann MM, Radke JB, Lucas O, Nawas J, Sibley LD, White MW. 2010. Coordinated progression through two subtranscriptomes underlies the tachyzoite cycle of Toxoplasma gondii. PLoS One 5: e12354.

22. Bozdech Z, Llinas M, Pulliam BL, Wong ED, Zhu J, DeRisi JL. 2003. The transcriptome of the intraerythrocytic developmental cycle of Plasmodium falciparum. PLoS Biol 1:E5.

23. Qu Z, Weiss JN, MacLellan WR. 2003. Regulation of the mammalian cell cycle: a model of the G1-to-S transition. Am J Physiol Cell Physiol 284:C349–64.

24. Cross FR, Buchler NE, Skotheim JM. 2011. Evolution of networks and sequences in eukaryotic cell cycle control. Philos Trans R Soc Lond B Biol Sci 366: 3532–44.

25. Silmon de Monerri NC, Yakubu RR, Chen AL, Bradley PJ, Nieves E, Weiss LM, Kim K. 2015. The Ubiquitin Proteome of Toxoplasma gondii Reveals Roles for Protein Ubiquitination in Cell-Cycle Transitions. Cell Host Microbe 18: 621–33.

26. Li A, Blow JJ. 2005. Cdt1 downregulation by proteolysis and geminin inhibition prevents DNA re-replication in Xenopus. EMBO J 24: 395–404.

27. Li A, Blow JJ. 2004. Negative regulation of geminin by CDK-dependent ubiquitination controls replication licensing. Cell Cycle 3: 443–5.

28. Li A, Blow JJ. 2004. Non-proteolytic inactivation of geminin requires CDK-dependent ubiquitination. Nat Cell Biol 6: 260–7.

29. Tachibana KE, Gonzalez MA, Guarguaglini G, Nigg EA, Laskey RA. 2005. Depletion of licensing inhibitor geminin causes centrosome overduplication and mitotic defects. EMBO Rep 6: 1052–7.

30. Chen CT, Kelly M, Leon J, Nwagbara B, Ebbert P, Ferguson DJ, Lowery LA, Morrissette N, Gubbels MJ. 2015. Compartmentalized Toxoplasma EB1 bundles spindle microtubules to secure accurate chromosome segregation. Mol Biol Cell 26: 4562–76.

31. Francia ME, Dubremetz JF, Morrissette NS. 2015. Basal body structure and composition in the apicomplexans Toxoplasma and Plasmodium. Cilia 5: 3.

32. Ruthnick D, Schiebel E. 2016. Duplication of the Yeast Spindle Pole Body Once per Cell Cycle. Mol Cell Biol 36: 1324–31.

33. Dubremetz JF. 1973. [Ultrastructural study of schizogonic mitosis in the coccidian, Eimeria necatrix (Johnson 1930)]. J Ultrastruct Res 42: 354–76.

34. Ferguson DJ, Sahoo N, Pinches RA, Bumstead JM, Tomley FM, Gubbels MJ. 2008. MORN1 has a conserved role in asexual and sexual development across the apicomplexa. Eukaryot Cell 7: 698–711.

35. Sibert GJ, Speer CA. 1981. Fine structure of nuclear division and microgametogony of Eimeria nieschulzi Dieben, 1924. Z Parasitenkd 66: 179–89.

36. Decottignies A, Zarzov P, Nurse P. 2001. In vivo localisation of fission yeast cyclin-dependent kinase cdc2p and cyclin B cdc13p during mitosis and meiosis. J Cell Sci 114: 2627–40.

37. Alfa CE, Ducommun B, Beach D, Hyams JS. 1990. Distinct nuclear and spindle pole body population of cyclin-cdc2 in fission yeast. Nature 347: 680–2.

38. Krapp A, Schmidt S, Cano E, Simanis V. 2001. S. pombe cdc11p, together with sid4p, provides an anchor for septation initiation network proteins on the spindle pole body. Curr Biol 11: 1559–68.

39. Jaspersen SL, Winey M. 2004. The budding yeast spindle pole body: structure, duplication, and function. Annu Rev Cell Dev Biol 20: 1–28.

40. Roos DS, Donald RG, Morrissette NS, Moulton AL. 1994. Molecular tools for genetic dissection of the protozoan parasite Toxoplasma gondii. Methods in cell biology 45: 27–63.

41. Donald RG, Roos DS. 1998. Gene knock-outs and allelic replacements in Toxoplasma gondii: HXGPRT as a selectable marker for hit-and-run mutagenesis. Mol Biochem Parasitol 91: 295–305.

42. Huynh MH, Carruthers VB. 2009. Tagging of endogenous genes in a Toxoplasma gondii strain lacking Ku80. Eukaryot Cell 8: 530–9.

43. Gaji RY, Behnke MS, Lehmann MM, White MW, Carruthers VB. 2011. Cell cycle-dependent, intercellular transmission of Toxoplasma gondii is accompanied by marked changes in parasite gene expression. Molecular microbiology 79: 192–204.

44. Dereeper A, Guignon V, Blanc G, Audic S, Buffet S, Chevenet F, Dufayard JF, Guindon S, Lefort V, Lescot M, Claverie JM, Gascuel O. 2008. Phylogeny.fr: robust phylogenetic analysis for the non-specialist. Nucleic Acids Res 36:W465–9.

